# Analysis of DYRK1B, PPARG, and CEBPB Expression Patterns in Adipose-Derived Stem Cells from Patients carrying DYRK1B R102C and Healthy Individuals During Adipogenesis

**DOI:** 10.1101/2021.10.08.463642

**Authors:** Azam Armanmehr, Hossein Jafari Khamirani, Sina Zoghi, Mehdi Dianatpour

## Abstract

**Background:** Metabolic syndrome (MetS) is a group of signs and symptoms that are associated with higher risk of Type 2 Diabetes Mellitus (T2DM) and Cardiovascular Diseases (CVDs). The major risk factor for developing MetS is abdominal obesity that is caused by increase in adipocyte size or number. Adipocyte number multiplication is caused by differentiation of mesenchymal stem cells into adipose tissue. Numerous studies have evaluated the expression of key transcription factors including PPARG and CEBPB during adipocyte differentiation in murine cells such as 3T3-L1 cell line. In order to comprehend the expression changes during the process of fat accumulation in adipose tissue derived stem cells (ASCs), we compared the expression of DYRK1B, PPARG, and CEBPB in undifferentiated and differentiated ASCs into mature adipocytes between the patient (harboring DYRK1b R102C) and control (healthy individuals) groups.

**Methods:** Gene expression was evaluated on eighth days pre-induction and day 1, 5 and 15 post-induction. The pluripotent capacity of ASCs and the potential for differentiation into adipocyte were confirmed by flow cytometry analysis of surface markers (CD34, CD44, CD105 and CD90), and Oil red O staining, respectively. Expression of DYRK1B, PPARG, and CEBPB were assessed by RT-PCR in patients’ and normal individuals’ samples.

**Results:** The expression of DYRK1B kinase and transcription factors (CEBPB and PPARG) are significantly higher in adipose derived stem cells harboring DYRK1b R102C compared to non-carriers on day 5 and 15 during adipocyte differentiation. These proteins may be suitable targets for therapeutic strategies in obesity and obesity related disorders like metabolic syndrome. Furthermore, AZ191 exhibited a potent and selectively inhibitory activity toward DYRK1B and CEBPB.

**Conclusion:** CEBPB, PPARA and DYRK1B contribute to adipogenesis and the development of metabolic syndrome; thus, they can be harnessed in developing therapeutic agents against metabolic syndrome.

## Introduction

Metabolic syndrome (MetS) or X syndrome is a group of signs and symptoms comprising abdominal obesity, elevated blood pressure, dyslipidemia and insulin resistance(Kaur, 2014). MetS is roughly related to two-fold increase in the risk of Cardiovascular Diseases (CVDs) and five-fold increase in the risk of Type 2 Diabetes Mellitus (T2DM)(Azhar, 2010; Martin et al., 2015). Depending on various criteria including place of residence, ethnicity, and the definition used to diagnosis, the prevalence of MetS is estimated to be ranging from <10% to as high as 84% around the world(Desroches and Lamarche, 2007; Kolovou et al., 2007). The prevalence of this syndrome is relatively high in Iran, with the estimates ranging from 20% to 25%(Amirkalali et al., 2015). Several variants in DYRK1B has been reported. Among these variants, NM_004714.3: p.Arg102Cys and NM_004714.3: p.His90Pro are linked to metabolic syndrome with autosomal dominant inheritance pattern(Keramati et al., 2014). The DYRKs (dual-specificity tyrosine-phosphorylated kinases) is a highly conserved family of protein kinases. In mammalian DYRKs are divided into two classes. Class I (DYRK1A and DYRK1B) and class II (DYRK2, DYRK3, and DYRK4)(Aranda et al., 2011) These proteins participate in many vital processes such as metabolism, gene expression and signaling (Lahiry et al., 2010). As dysregulation of these kinases are linked to multiple conditions like diabetes and CVD, they have been considered as powerful target for treatment of MetS(Fabbro, 2015; Ghoreschi et al., 2009; Masaki et al., 2015). DYRK1B participates in adipocyte differentiation and undergoes various splicing during fat formation(Ashford et al., 2016). There is increasing efforts and investigations in designing effective inhibitors of DYRKs. INDY and Harmin show inhibitory effect on DYRK1A and DYRK1B(Göckler et al., 2009; Ogawa et al., 2010). AZ191, a new compound, is a selective inhibitor of DYRK1B. AZ191 like harmine selectively inhibits Ser/Thr kinase activity at 1µ M concentration(Ashford et al., 2016).

**Figure 1.**
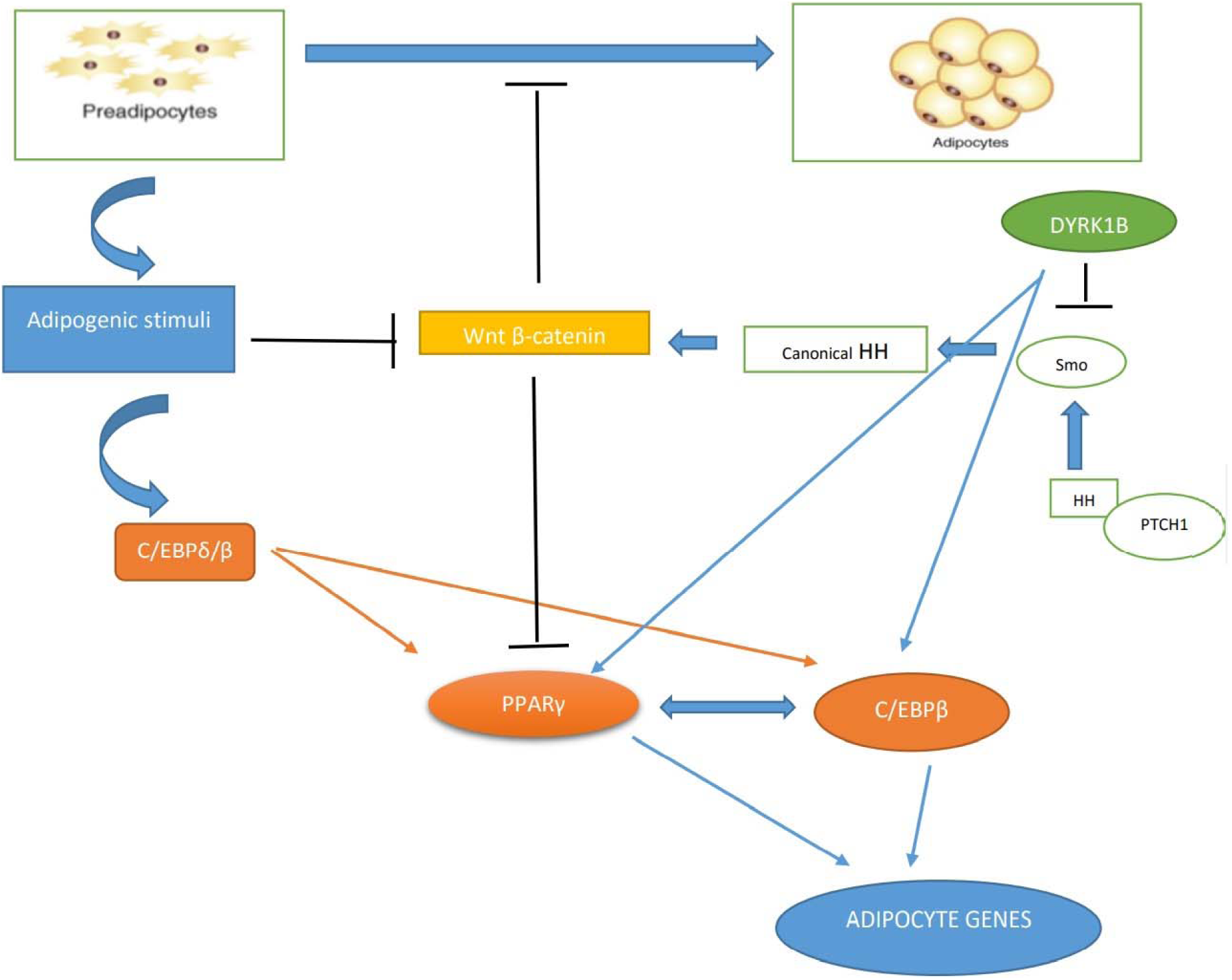
Summary of the function of DYRK1B, CEBPDB, CEBPB and PPARG during adipogenesis.

It has been illustrated that R102C promotes adipogenesis and causes MetS by increasing obesity(Foroozanfar et al., 2015; Keramati et al., 2014; Masaki et al., 2015). Accumulation of adipose tissue results in obesity(Cousin et al., 2007). Adipocyte differentiation is a well-organized multistep process requiring the successive activation of several groups of transcription factors, including CCAAT/enhancer-binding protein (CEBP) family and peroxisome proliferator-activated receptor-γ (PPARG). During adipocyte differentiation, CEBPB is expressed early and induces the expression of CEBPγ and PPARG, two master transcription factors for terminal adipocyte differentiation. Numerous studies have utilized pre-adipocyte cell lines like 3T3-L1 or 3T3-442A to evaluate molecular mechanisms affecting adipogenesis.. The culture medium contains the enhancers of adipogenesis IBMX and dexamethasone, which regulate PPARG. Furthermore, IBMX and dexamethasone are stimulators of transcription factors CEBPB⍰ and CEBPL which are essential for differentiation(Scott et al., 2011). In the present study, we focus on DYRK1B, PPARG, and CEBPB expression alterations in adipose tissue derived stem cells harboring p.Arg102Cys and compare them with non-affected cells on days 1, 5, and 15 during adipogenesis.

**Figure 2.**
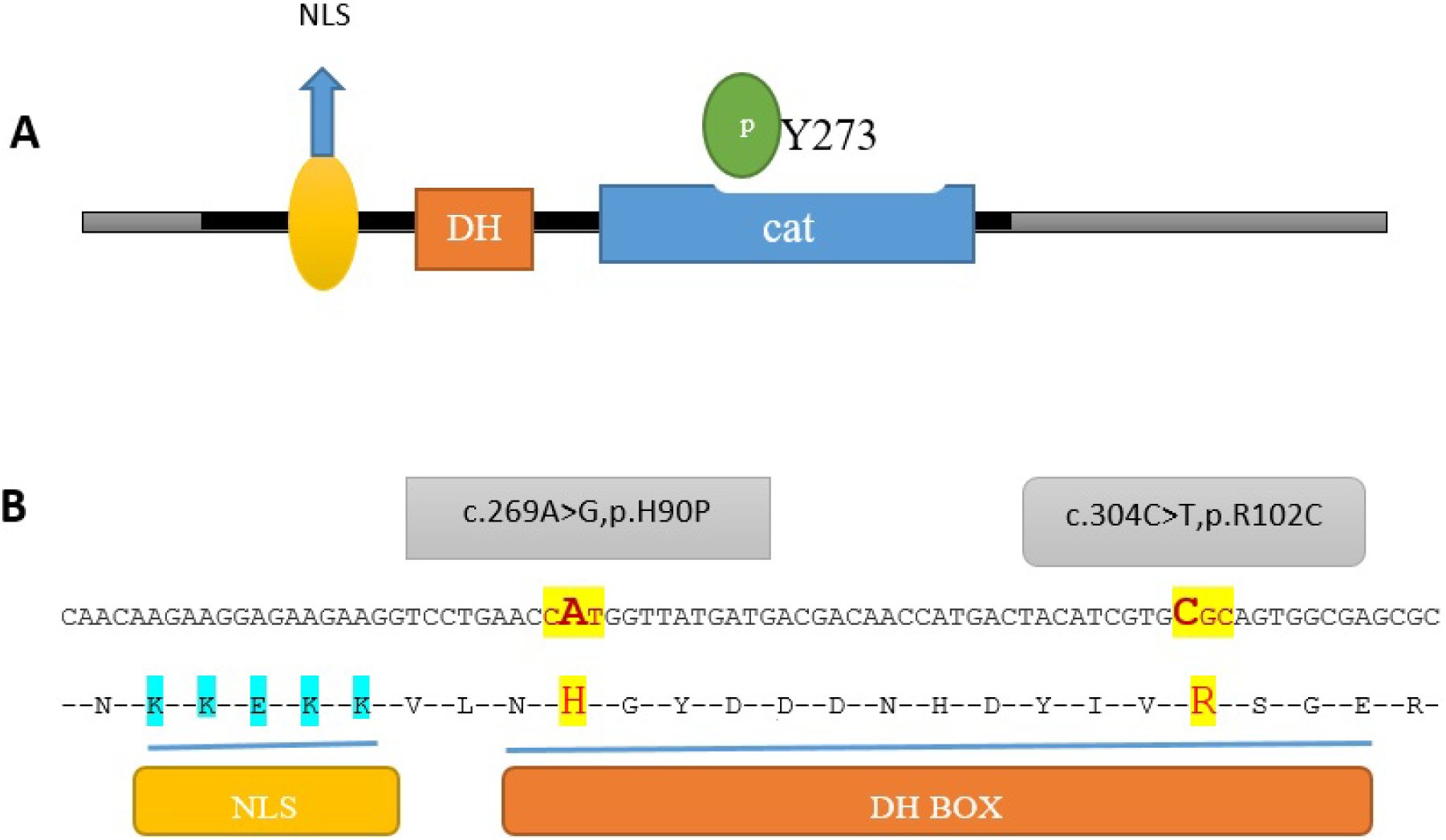
A Domain structure of DYRK1B. The location of DH box is between the nuclear localization signal (NLS) and the catalytic domain (cat).The autophosphorylation of the tyrosine (pY273) is shown by green circled P in the activation loop. B The position of H90P and R102C in DH box.

## Material and Methods

### Adipose Stromal Cells Culture, Immunophenotypic Characterization and Induction of Adipogenic Differentiation

Subcutaneous adipose tissue was obtained from three adult patients harboring the variant and three adult normal individuals (including four male and two female individuals) under local anesthesia by a board-certified surgeon. Cells from the affected and unaffected participants were cultured in complete culture medium encompassing DMEM (GIBCO, Life Technologies™, New York, USA) complemented with 10% fetal bovine serum (FBS; Sigma-Aldrich, St. Louis, MO, USA), 1% (v/v) penicillin [10,000 U/mL]–streptomycin [10,000 μg/mL] (p/s; GIBCO, Life Technologies™, New York, USA), and 1% non-essential amino acids in 75 cm2 culture flasks (NUNC™, Roskilde Site, Kamstrupvej, Denmark) under humidified standard condition at 37 °C in 5% CO2 atmosphere (Forma water jacketed CO2 incubator, 3111TF; Thermo Fisher Scientific, Waltham, Massachusetts). When the cells reached 80% confluence, they were washed twice with 3 mL Phosphate Buffer Saline (PBS) containing 2% (v/v) p/s and separated from the bottom of culture flask by adding 1 mL trypsin (GIBCO, Life Technologies™, New York, USA) then incubated at 37 °C in 5% CO2 for about 7–10 minutes to ensure that the cells were separated from the flask walls. Trypsin was deactivated by adding the same amount of complete medium. The suspension was centrifuged at 1,200 rpm for 5 minutes to pellet the cells.

### Surface Marker Characterization

The MSCs were characterized by the analysis of expression of cell surface markers with flow cytometry at the third passage. The cell suspension (1×106 cells/mL) was washed in blocking solution, cold PBS containing 10% FBS, for 20 minutes. Subsequently, the cells were labeled with Fluorescein isothiocyanate (FITC)-conjugated anti-CD44, anti-CD90, Phycoerythrin-conjugated anti CD34, and PerCP (Peridinin Chlorophyll Protein complex)-conjugated anti-CD105 antibodies (Abcam, Cambridge, UK). They were then washed with cold PBS, resuspended in PBS containing 10% FBS, and analyzed by flow cytometery (FACS Calibur™, BD Biosciences, San Jose, CA). The data were analysed by utilizing the WinMDI 2.9 software.

### Adipogenic Differentiation

For adipogenic differentiation, cells were seeded at a density of 5000 cells/cm2 in complete culture medium and grown to confluence. The cell cultures were washed twice with 3 mL PBS containing 2% (v/v) p/s and differentiated by adding adipogenic induction medium to the cell cultures. The adipogenic medium contained Dulbecco’s Modified Eagle’s Medium (GIBCO, Life Technologies™, New York, USA) completed with 10% FBS, 0.5Mm isobutyl-methylxanthin (IBMX), 1μM dexamethasone, 10 μM insulin, 200 μM indomethacin (Sigma-Aldrich, St. Louis, MO, USA). Cells from passages 3-5 were used. The process of differentiation was completed during a 15-day period. Cells were harvested on three different time points including days 1, 5, and 15 and used for RNA isolation.

### Oil Red O staining

The completion of adipogenesis was confirmed by Oil Red O, which stains intracellular triglyceride droplets. After 15 days of differentiation, the cells were fixed with 10% formalin and then incubated for 15 minutes with Oil Red O solution. Thereafter, the cells were washed three times with distilled water and the dye was eluted from the cells using isopropanol.

### RNA isolation and RT-qPCR

Total cellular RNA was isolated from the control and patient differentiated cells by RNA extraction kit (Cinnagen Inc., Tehran, Iran). The quantity and quality of obtained RNA was checked using NanodropTM spectrophotometer (3411 Silverside Rd, Bancroft Building Wilmington DE, 19810. USA) and agarose gel electrophoresis, respectively. Samples were stored at -80°C until cDNA synthesis. The cDNA was synthesized using 500 ng of total RNA in a first-strand complementary DNA synthesis reaction by the help of RevertAid™ First Strand cDNA Synthesis kit (Fermentas Inc., Schwerte, Germany). Quantitative real-time PCR was performed using the 7500 Real Time PCR System and the RealQ Plus 2x Master Mix Green (Ampliqon Co., Denmark). Specific primers targeting DYRK1B, PPAR-γ and CEBPB were designed using Allel ID (version 7.5) (Table 1). 200 nM of each primer (Table 1) was added into each reaction to target the specific sequence. The GAPDH housekeeping gene was also used as internal control of qPCR reactions. The qPCR was set at 15 minutes at 94°C followed by 45 cycles of 30 seconds at 94°C, 1 minute at 60 °C.

**Table1.**
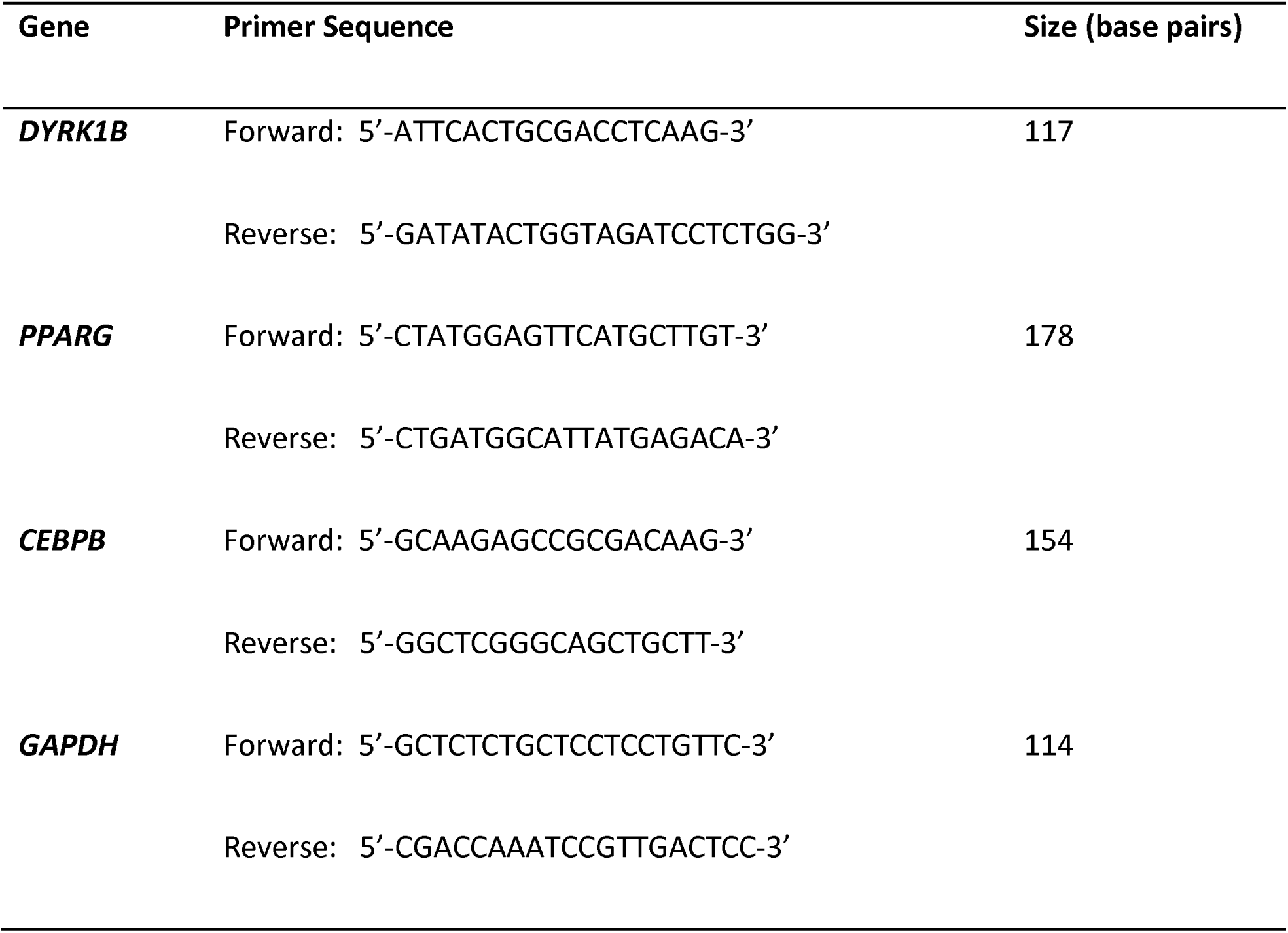
Real-time PCR primers used in this study

We performed all in vitro experiments in two groups (including 3 normal and 3 patient individuals). GraphPad software (2365 Northside Dr. Suite 560 San Diego, CA 92108) was used to illustrate the data.

### Statistical analysis

Statistical comparison was performed using independent-sample t-test.

All individuals participating in this study are from a family living in southwestern Iran. Written informed consent for participation and publication was obtained from all of the individuals participating in this study, individually. The Ethnic Committee of Shiraz University of Medical Sciences, Iran approved the present study (Ethics Code: IR. SUMS. REC. 1395.5604).

## Results

### Differentiation of AMSCs

The adipogenic differentiation of MSCs was confirmed by Oil Red O staining (Figure 3). In the presence of adipogenic media, MSCs could store lipid droplet. These results confirmed the pluripotent capacity of MSCs(Ghorbani et al., 2014).

**Figure 3.**
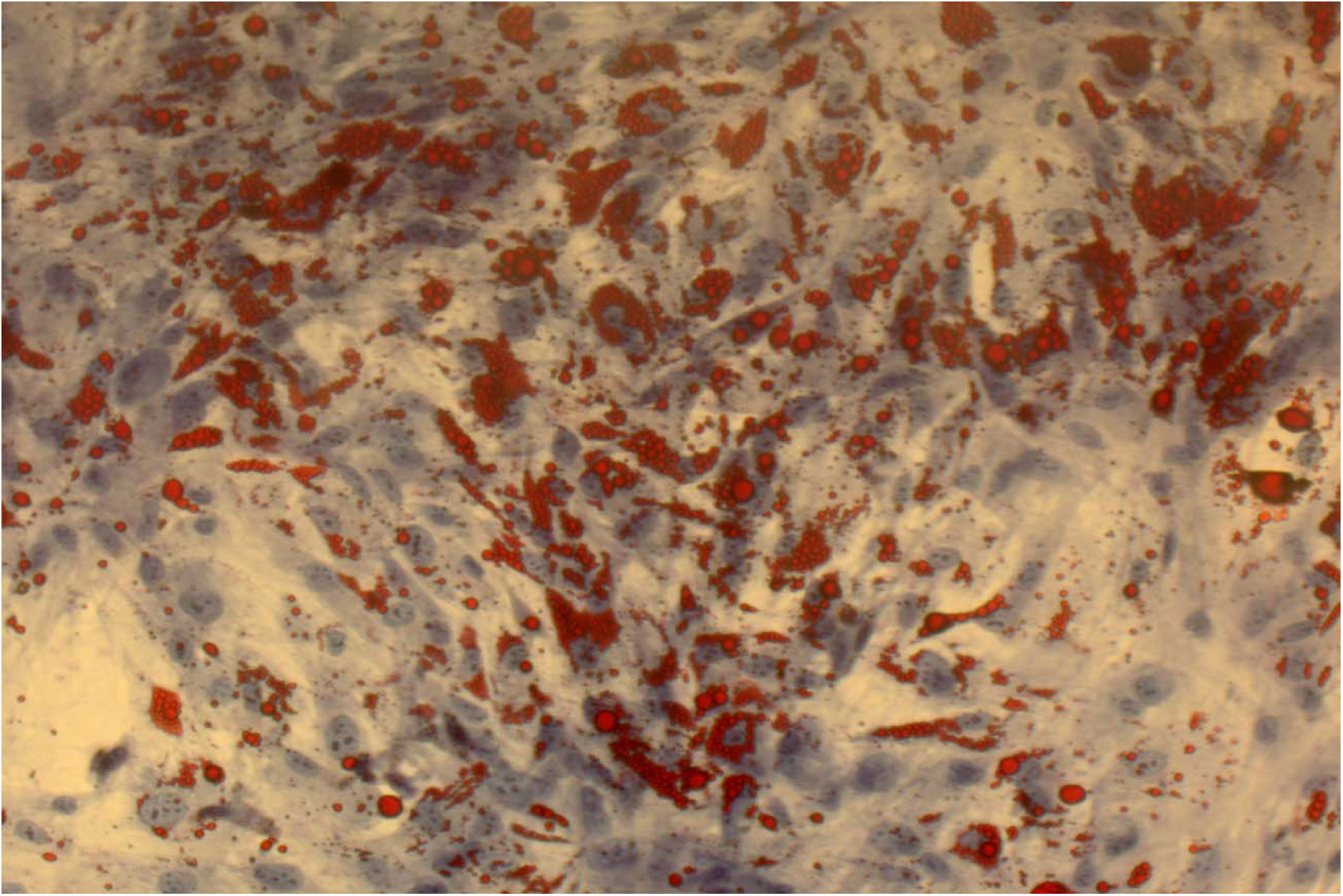
Stored lipid droplets in the presence of adipogenic medium stained with Oil Red O on day 10 during adipogenesis.

### Quantitative real-time PCR (qRT-PCR)

The DYRK1B, PPARG and CEBPB were upregulated on days 5 and 15, indicating the maturation of the adipocytes. The expression of DYRK1B, PPARG and CEBPB were increased on 5th and 15th day in comparison with the first day since initiation of differentiation.

The patients harboring the variant had an overall higher rate of expression of the DYRK1B, PPARG and CEBPB in comparison with the controls. The expression of PPARG and CEBPB was highest on the 15th day in the control. The expression patterns of DYRK1B, PPARG and CEBPB in control and the patients population are summarized in Figure 4.

**Figure 4.**
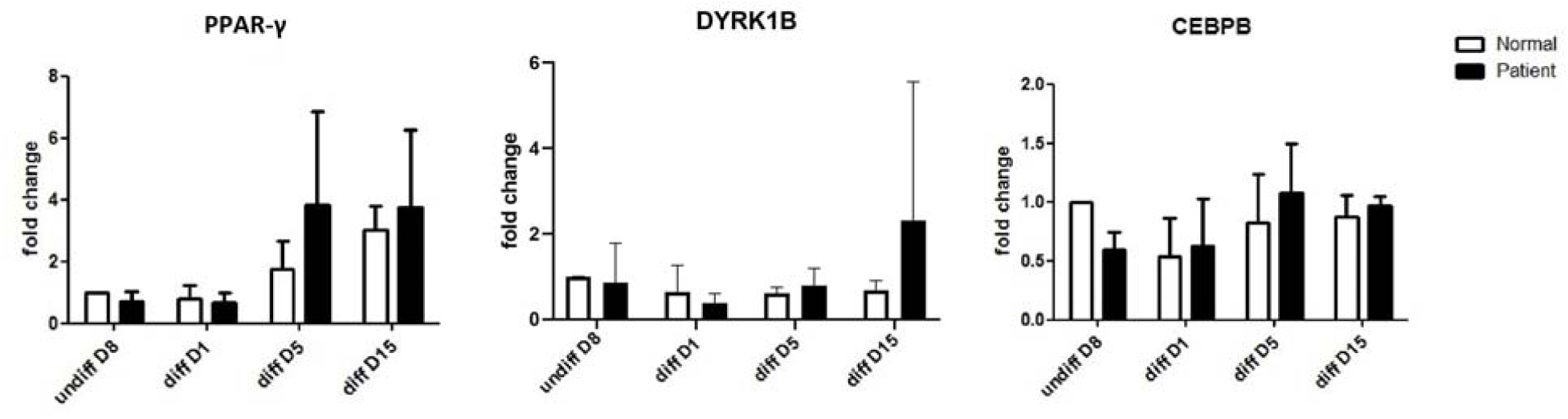
The expression levels of DYRK1B, PPAR-γ, and CEBPB in samples obtained from patents and normal individuals.

### The Effect of AZ191 on the Expression of DYRK1B and CEBPB

The Effect of AZ191 on the Expression of DYRK1B and CEBPB was assessed in the complete culture medium (control) and adipogenic mediua with and without AZ191 at 3µ M concentratin. The expression of DYRK1B and CEBPB were lower in the medium enriched by AZ191 (as shown in Figure 5).

**Figure 5.**
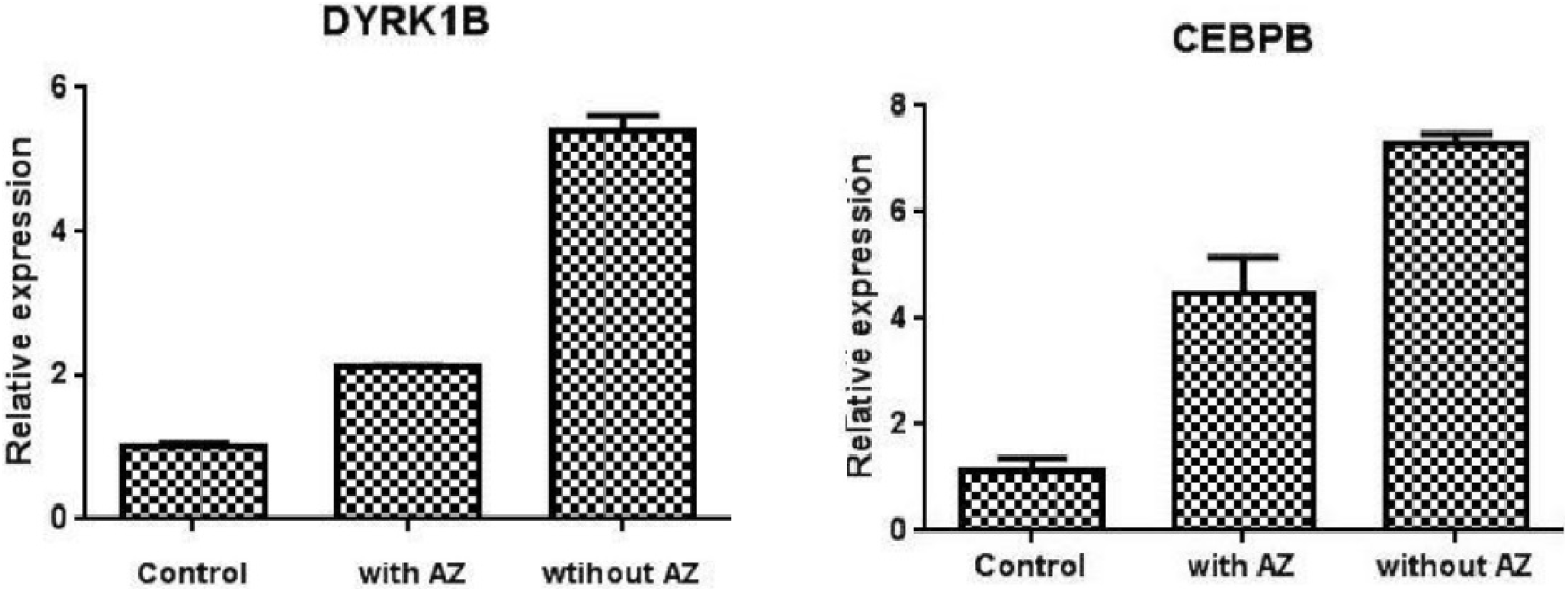
The expression of DYRK1B and CEBPB in various media in the cells harboring the variant in the tenth day of differentiation.

### Identification of Cell Surface Markers

The identification of was based on the presence of CD44, CD105, and CD90 and absence of markers specific to hematopoietic cell line such as CD34. As shown in Figure 6, the expression of CD34 was as low as 0.7% and CD44, CD105, and CD90 were detected on the surface of 94.6%, 52.6%, and 96.9% of cells. These results are comparable with other studies and confirm the existence of mesenchymal stem cells(Dominici et al., 2006; Wongchuensoontorn et al., 2009; Zuk et al., 2002).

**Figure 6.**
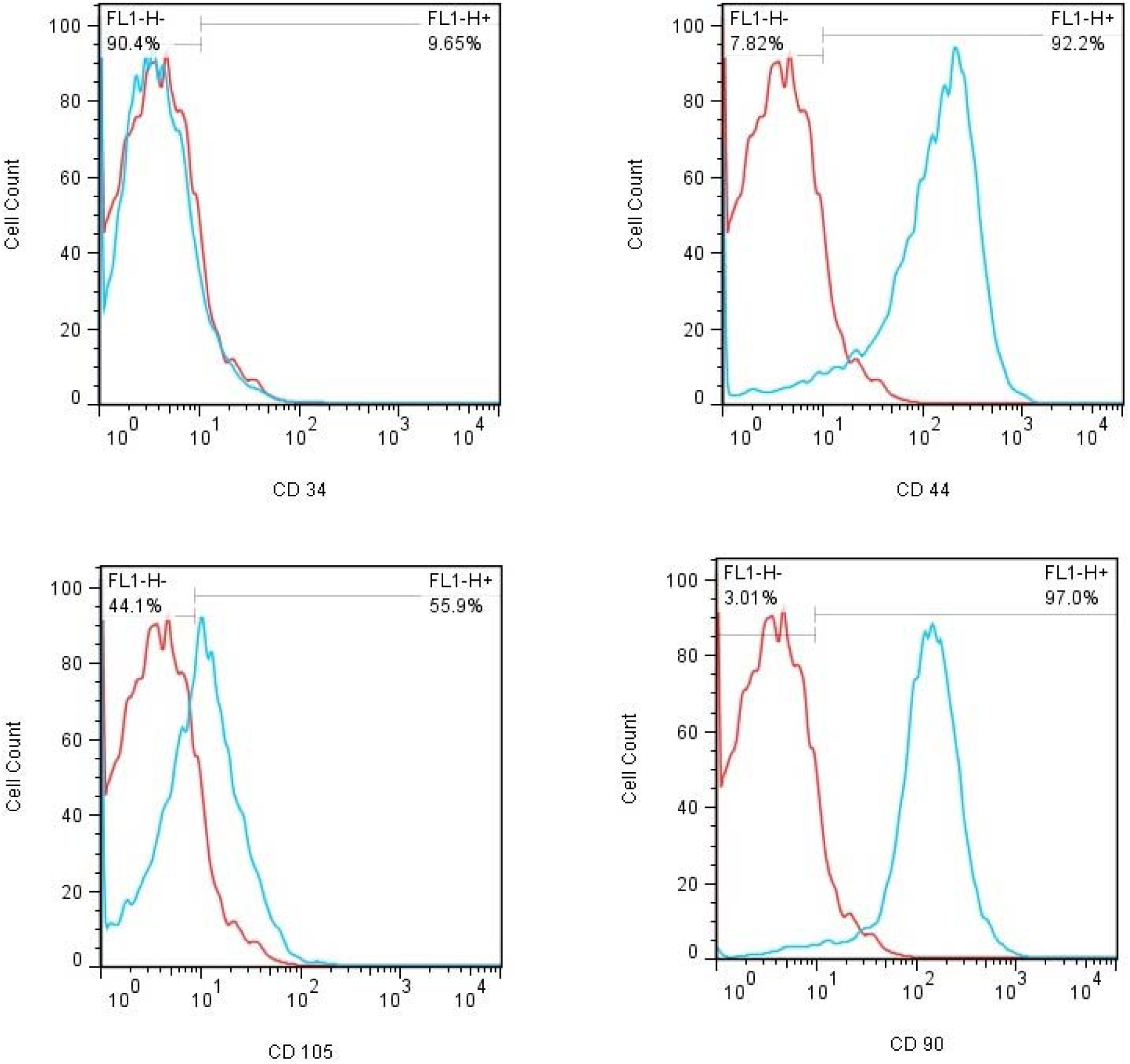
Histograms resulting from flow cytometry showing the immunophenotype of the adipose tissue-derived mesenchymal stem cell.

## Discussion

DYRK1B is a member of DYRK family, comprising evolutionary conserved protein kinases(Azhar, 2010; Kaur, 2014). DYRK1B is involved in several processes including myogenesis, adipogenesis, carcinogenesis, and cell survival(Azhar, 2010). In comparison to murine cell lines, human mesenchymal stem cells are more troublesome to manipulate, nevertheless their usefulness and unique curative potential makes them necessary to study. Moreover, differences in basic structure and regulation between human and rodents may affect gene expression and, as a result, murine cells would be less reliable to model human diseases like obesity(López et al., 2003).

Presently, the literature regarding the role of CEBPB, PPARG, and DYRK1B are focused on studies on animals and in vitro investigations in human are quite rare. Previous studies showed that DYRK1B R102C enhances adipogenic differentiation(Haldar, 2014).Anti-adipogenic characteristic of Hedgehog signaling could increase the expression of their anti-adipogenic target genes like Wnt. Activation of Wnt protein family hinders adipocyte differentiation (Cousin et al., 2007). Wnt signaling restrains adipocyte differentiation by inhibiting the expression of key adipogenic transcription factors, namely PPARG and CEBPA. The expression of these transcription factors are induced by CEBPB and CEBPD(Christodoulides et al., 2009). It has been demonstrated that DYRK1B stimulates adipogenesis by inhibiting SHH pathway(Keramati et al., 2014).

Mutated DYRK1B (p.Arg102Cys) was previously reported to have reduced activity(Ashford et al., 2016). Nevertheless, it was shown that p.Arg102Cys is not directly involved in the function of DYRK1B. This residue is not evolutionarily conserved in DYRK1B orthologs, like other amino acids that directly participate in enzyme function. This missense mutation renders DYRK1B vulnerable to misfolding and degradation or aggregation and, as a result, confers a loss of biochemical function rather than a gain of function. Although loss-of-function mutations are usually observed to inherit in a recessive fashion, it plausible that the mutated DYRK1B aggregation causes a gain of toxic function(Jhaisha et al., 2017).

Based on our information, this is the first study evaluating the possible role of CEBPB, PPARG DYR1B in the development and progression of the metabolic syndrome. The genes that are expressed during the late phase of adipocyte differentiation may play major role in promoting excessive proliferation and accumulation of lipid droplets, which contribute to the development of obesity and obesity-related disease. Previous studies indicate that gain-of-function and loss-of-function mutation in PPARG lead to obesity and decreased body weight, respectively(Fei, 2012). Besides PPARA, there is strong evidence that CEBPB plays an important role in adipocyte differentiation, suggesting implications in obesity(van der Krieken et al., 2015). Wnt and Hedgehog signaling inhibit adipocyte differentiation via affecting PPARG, a key regulator of adipogenesis(Moseti et al., 2016). Previous studies have shown that the increased level of CEBPB result in inhibiting Wnt signaling. Moreover, CEBPB is capable of activating adipocyte maturation by inducing PPARA which in turn, leads to activating CEBPG (van der Krieken et al., 2015). In an agreement with these studies, PPARG and CEBPB are significantly highly expressed during adipogenesis, confirming their prominent role in adipogenesis.

In this study, we have shown the importance of NM_004714.3: p.Arg102Cys in altering the expression levels of DYRK1B, PPARG, and CEBPB, which underlies the obesity and subsequent metabolic syndrome in the patients harboring this variant. This study on top of shedding light on the effect of the variant on the expression of DYRK1B, PPAR-γ, and CEBPB, shows the in vitro efficacy of AZ191 in downregulating the increased expression of the aforementioned proteins, possibly counteracting the effects of the variant and the subsequent obesity it causes. Moreover, new biological treatments including cell therapy, gene therapy, iRNA, etc. can be utilized to treat those suffering from metabolic syndrome caused by this variant(Gao and Liu, 2014).

## Conclusions

In conclusion, the potential role of CEBPB, PPARA and DYRK1B in adipogenesis indicate their contribution to the development of metabolic syndrome. Furthermore, it has not been clear that whether DYRK1B variants cause metabolic syndrome by inducing adipogenesis or through glucose production. Thus clarifying the molecular mechanism of DYRK1B mutations and functional studies of these transcriptional factors is essential.

## Declarations

### Ethical approval and consent to participants

Written informed consent was obtained from each patient or, in the case of minors, from their parents. This study was approved by The Ethics Committee of Shiraz University of Medical Sciences.

### Consent to publish

Consent for participation and publication was obtained from the patients or, in the case of minors, from their parents.

### Availability of Data and Material

All data generated or analyzed during this study are included in the final published article.

### Authors’ contributions

This study was designed and conceptualized by AA and MD. AA, HJK, and SZ executed the study. MD supervised the execution of the study. AA drafted the manuscript and SZ revised the final version. All authors approved the final version of the manuscript.

